# AUK3 is required for faithful nuclear segregation in the bloodstream form of *Trypanosoma brucei*

**DOI:** 10.1101/2024.09.24.614706

**Authors:** J.A. Black, B.C. Poulton, B.S. Gonzaga, A. Iskantar, L.R.O. Tosi, R McCulloch

**Author notes:** Corresponding Author: Richard McCulloch.

## Abstract

Eukaryotic chromosomes segregate faithfully prior to nuclear division to ensure genome stability. If segregation becomes defective, the chromosome copy number of the cell may alter leading to aneuploidy and/or polyploidy, both common hallmarks of cancers. In eukaryotes, aurora kinases regulate chromosome segregation during mitosis and meiosis, but their functions in the divergent, single-celled eukaryotic pathogen *Trypanosoma brucei* are less understood. Here, we focused on one of three aurora kinases in these parasites, TbAUK3, a homologue of the human aurora kinase AURKC, whose functions are primarily restricted to meiosis. We show that RNAi targeted depletion of TbAUK3 correlates with nuclear segregation defects, reduced proliferation, and decreased DNA synthesis, suggestive of a role for TbAUK3 during mitotic, not meiotic, chromosome segregation. Moreover, we uncover a putative role for TbAUK3 during the parasite’s response to DNA damage since we show that depletion of TbAUK3 enhances DNA instability and sensitivity to genotoxic agents.

**HIGHLIGHTS:**

- The C-terminus of TbAUK3 is disordered
- TbAUK3 depletion coincides with nuclear segregation defects
- Depletion of TbAUK3 enhances DNA instability

## MAIN ARTICLE

*Trypanosoma brucei* (*class* Kinetoplastea) are pathogenic protozoans of medical importance, causing the Neglected Tropical Disease Human African Trypanosomiasis (HAT) in humans, or Animal African Trypanosomiasis (AAT; ‘Nagana’) in livestock [1, 2]. Kinetoplastids display considerable divergence from more commonly studied eukaryotes in several key cellular processes and possess advanced molecular genetic tools, meaning they offer a window into the diversity of protist and eukaryotic biology.

Eukaryotic chromosomes segregate by connecting spindle microtubules and centromeric DNA via macromolecular complexes called kinetochores, whose subunits bear very little, if any, homology between *T. brucei* and model eukaryotes like yeast [3]. If correctly assembled, the spindle assembly checkpoint (SAC) is lifted, and chromosome segregation begins [4]. Numerous factors orchestrate this process, including aurora kinases, of which mammalian cells express three: AURKA, AURKB and AURKC. Functions of AURKA and AURKB are largely linked to mitosis, whereas AURKC primarily functions during meiosis, with expression restricted to meiotically-active germ line cells (spermatozoa and oocytes) and peaking during meiosis [5, 6].

Here, AURKC acts as a component of the chromosomal passenger complex (CPC), regulating the meiotic SAC. Mutations of AURKC correlate with male infertility [7–10]. Expression of AURKC is typically low in somatic cells, with higher expression levels detected in some cancers (e.g., breast cancer); specifically, enhanced AURKC expression coincides with supernumerary chromosomes and appearance of polyploid cells [11–13]. Functions of the SAC and chromosome segregation during mitosis and meiosis are well described in model eukaryotes, whereas *T. brucei* likely lacks a canonical SAC [14]. This is evidenced by; 1) the absence of SAC factors in their genomes, except for TbMad2 [15] and, more recently, Bub1-like/BubR1-Bub3 like kinetochore proteins, KKT14-KKT15 [16, 17]; 2) distinct subcellular localization patterns for TbMad2 and the trypanosome kinetochore complex; and 3) failure to correctly divide the nucleus does not block entry to cytokinesis [18]. Unexpectedly, TbMad2 localizes to the basal body (BB) of the parasite’s single flagellum, suggesting the BB may act in nuclear chromosome segregation [15]. Functional phenotyping of the three aurora kinases in *T. brucei* (TbAUK1, TbAUK2 and TbAUK3) has been largely restricted to TbAUK1 and TbAUK2. TbAUK1 orchestrates the mitotic metaphase-anaphase transition, thereby modulating chromosome segregation during mitosis and cytokinesis[16, 19–21], whereas TbAUK2 may have broader roles during DNA damage repair [22]. Data from a prior study implicated TbAUK3 in parasite proliferation and cell cycle regulation in bloodstream form cells (BSF; mammalian stage) [23]. More recently, in procyclic form cells (PCF; insect stage) TbAUK3 was localized to the BB, like TbMad2, and shown to exhibit a cell cycle-regulated localization pattern dissimilar to mammalian cells: TbAUK3 was detectable from late G1 until mid-to-late S-phase, during which time the kinetoplast DNA (K, mitochondrial), but not the nuclear DNA (N), completes segregation [15]. In a recent BSF RNAi screen TbAUK3 expression appears to peak around G2/M, though the emergence of a phenotype by RNAi may not entirely reflect the timing of expression for any gene factor [24]. However, this expression profile is similar to reports from mammalian cells [25, 26]. Taken together, these data present a potentially incomplete view of the role of TbAUK3 in *T. bruce*i, but perhaps argue against a function in meiosis, despite evidence of sexual exchange and meiotic proteins encoded in the genome [27, 28]. Here, we sought to understand the roles TbAUK3 in greater depth, with the hope of uncovering novel, parasite-specific biology or capturing information on Aurora kinase biological activities prior to *T. brucei’s* divergence from other eukaryotes.

TbAUK3 shares 39.85 % homology (2e^-61^; 83 % coverage) with the human kinase AURKC (HsAURKC), primarily within the kinase domain (**Fig. S1; Fig. S2A**). By aligning the amino acid sequence of TbAUK3 to HsAURCK we tested for key domains typically associated with kinase activity and aurora kinase biology (**Fig. S1**). Several key kinase-associated domains were identifiable in the sequence of TbAUK3, suggesting the protein is an active kinase (**Fig. S1; Fig. S2A**). We additionally noted conservation of the autophosphorylation site T198 (in AURKC; T246 in *T. brucei*), located within the activation loop of the kinase. HsAURKC has four D-box domains (‘RXXL’), which may play a role in protein degradation [6], whereas TbAUK3 has a single D-box domain, whose function has not been tested. In addition, we detected a DFS motif within the substrate binding loop of TbAUK3. Since most eukaryotic kinases (ePKs) possess a ‘DFG’ motif, with the glycine (G) allowing flexibility within the active site [29], the implications of a serine (S) within the region are unclear. However, recent characterisation of another *T. brucei* kinase revealed that the ‘DFS’ motif may enhance kinase stability, putatively locking the kinase in a more rigid ‘pre-activation’ state [30]; whether this holds true for TbAUK3 remains to be tested. When examining the AlphaFold2-predicted structure of TbAUK3 we noted the C-terminus of the kinase was disordered (**Fig. 1A**), and that this domain almost doubled the size of the parasite kinase relative to the human counterpart (HsAURKC, 309 AA; TbAUK3, 650 AA) (**Fig. 1B; Fig. S2B&C**); the function of this extension is unknown. In all, TbAUK3 shares features of a typical, active ePK in addition to divergences, some of which could confer parasite-specific functionality.

**Fig. 1.**
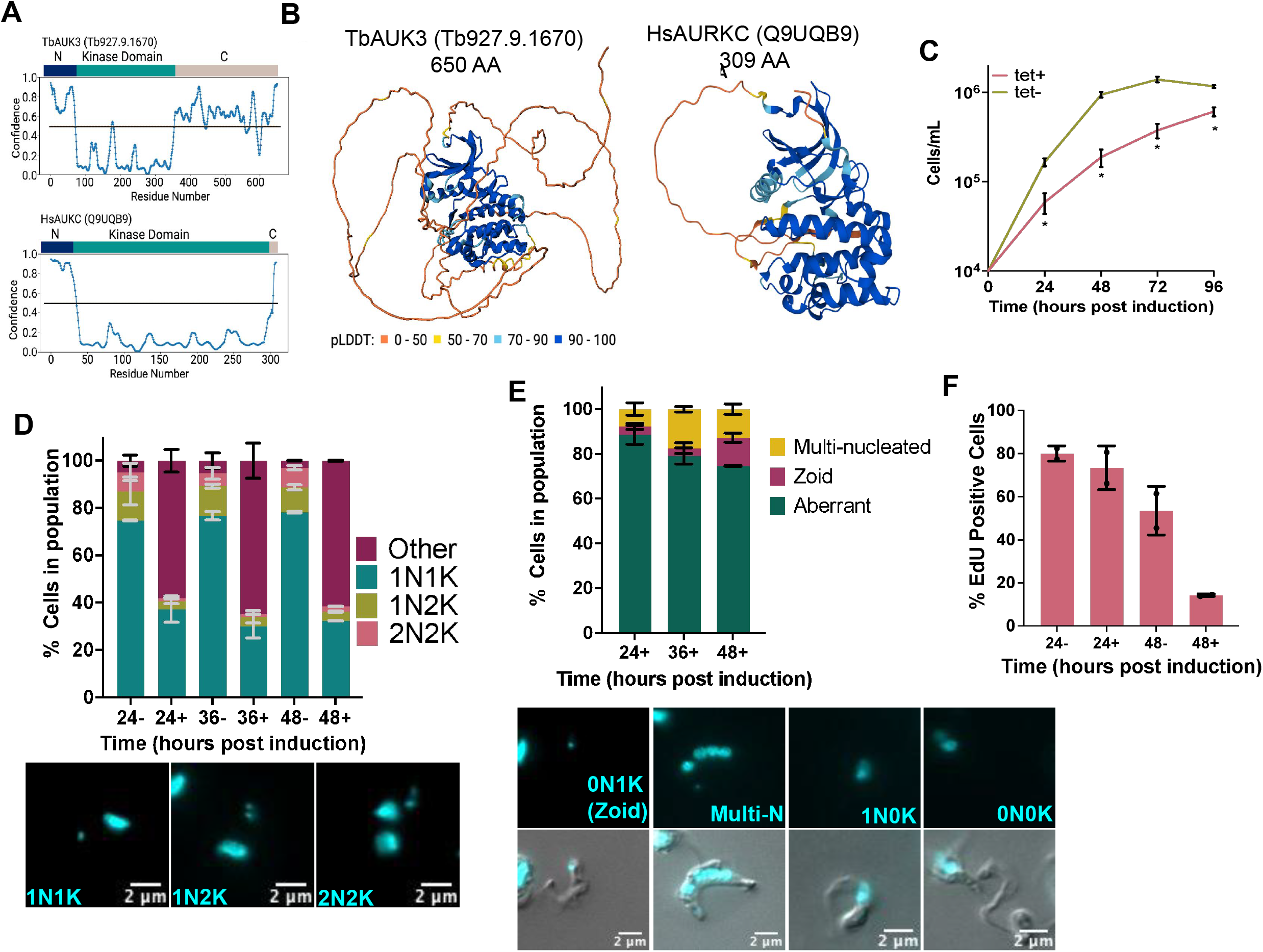
RNAi depletion of TbAUK3 in BSF *T. brucei* cells reduces parasite proliferation and correlates with aberrant nuclear division. (A) We used the PrDOS server to predict disordered regions of TbAUK3 (Tb927.9.1670) and HsAURKC (Q9UQB9) using their respective amino acid sequences. Prediction false positive rate (∼ 5 %) (B) Alphafold2 models of TbAUK3 and HsAURKC. The pLDDT score indicates the level of confidence for each residue called scaled from 0-100. Higher scores are associated with higher confidence and typically higher accuracy of the prediction (C) Growth curve analysis of TbAUK3 RNAi cells. Cells were seeded at 1 ×10^4^ cells. mL^-1^ then tet added to induce RNAi depletion of TbAUK3. Control cells were grown in the absence of tet. Cell density was assessed by counting on a Neubauer chamber every 24 hrs until 96 hrs post induction. Error bars = +/-SEM, Significant was calculated by paired t test * = > 0.005, n = 6 independent experiments (D) The number of cells in the population categories by their nuclear (N) and kDNA (K) configurations. Values are expressed as a percentage of the total. Error bars = +/-SD, n = 2 independent biological experiments, with > 130 cells counted each time point. Representative images of N:K configurations are shown in the image panel below. Nuclear and kDNA is stained using DAPI (cyan). (E) Graph shows the percentage of these aberrant configurations expressed as a total of the number of aberrant cells. Error bars = +/-SEM, n = 2 independent experiments Representative images of aberrant nuclear configurations after depletion of TbAUK3 at 24 hrs post induction are shown below. (F) The total number of cells in the population positive for nuclear EdU signal, expressed as a percentage of the total number of cells counted. Error bars = +/-SD, n = 2 independent experiments, > 100 cells were counted/experiment.

Previously, tetracycline (tet) inducible RNA interference (RNAi) targeting TbAUK3 in BSF cells induced an accumulation of morphologically aberrant cells (termed ‘others’) and reduced parasite proliferation [23]. We confirmed these findings by depleting TbAUK3 using RNAi in BSFs by adding 1 µg. mL^-1^ tet (prepared in 70 % ethanol), confirming depletion of the target mRNA by RT-qPCR (**Fig. S2D&E**). To assess proliferation *in vitro*, we seeded parasites at a density of 1× 10^4^ cells. mL^-1^ in fresh HMI-11 medium (10 % serum) then monitored parasite density every 24 hrs for 96 hrs by counting. As expected, TbAUK3 depletion significantly reduced parasite proliferation relative to uninduced controls (**Fig. 1C**). We next examined the cell cycle by immunofluorescence analysis (IFA) using DAPI staining to assess the cell cycle stage of individual cells. Parasites were fixed in 3.7 % paraformaldehyde, stained with DAPI, then cells were scored based on the ratio of kDNA (K) to nuclear DNA (N). In uninduced controls at 24, 36 and 48 hrs, we detected a similar distribution of cell cycle stages (**Fig. 1D**; representative K:N configurations are shown to the right), with ∼ 75 % of cells in G1 of the cell cycle (1N1K), ∼ 12 % of cells S/G2 and ∼ 7 % of cells in mitosis (anaphase). Approximately 4 % of cells did not display any of these N:K ratios and were therefore classed as ‘other’. In contrast, depletion of TbAUK3 by RNAi led to a reduction of all cell cycle stages, with > 50 % of cells in the population displaying aberrant N:K configurations (‘other’) at 24, 36 and 48 hrs post induction. When we examined the nuclear and kDNA configurations of these ‘other’ cells in more detail, we observed numerous configurations of nuclear and kDNA **(Fig. 1E**). Due to the range of configurations, we opted for three categories: 1) aberrant cells which included cells with no nuclear or kDNA content (0N0K); cells with no kDNA but nuclear DNA (1N0K); any cell that did not conform to any other definable category 2) zoid (1K0N); and 3) multinucleated cells (multi-N). At 24 hrs post induction (**Fig. 1E; Fig. S2F**), ∼12 % of the population were either zoid (∼4%) or multinucleated (∼ 8%) cells. The number of multinucleated cells increased at 36 (∼ 17 %) and 48 hrs (∼ 12 %). At 36 hrs the number zoids of appeared unchanged (∼ 3% of cells in the populations), while at at 48 hrs, this increased substantially to ∼ 13 % of the total population. We additionally confirmed by flow cytometry an increase in both sub-G1 and >2N populations after depletion of TbAUK3 (**Fig. S2G**). Taken together, these data suggest loss of TbAUK3 leads to widespread defects in mitotic nuclear segregation.

Roles are emerging for aurora kinases in DNA replication initiation [31], and therefore we asked if DNA synthesis was altered by the loss of TbAUK3. To do this, we grew *T. brucei* in Thymidine-free Iscove Modified Dulbecco Medium (IMDM) for 24 hrs, then induced RNAi. Four hours prior to each timepoint, the cells were pulsed with 150 µM 5-Ethynyl-2’-deoxyuridine (EdU; a thymidine analogue) then fixed in 3.7 % PFA. Using Click-IT^(R)^ technology, we fluorescently labeled the incorporated EdU and additionally stained with DAPI to visualize the nucleus and the kDNA (**Fig. 1F; Fig. S2H**). In the absence of RNAi induction, and after 24 and 48 hrs of RNAi induction, we counted the number of EdU positive parasites in the population. At 24 and 48 hrs of growth, we detected EdU signal in the nuclei of ∼ 80 % (24 hrs) and ∼54 % (48 hrs) of uninduced cells in the population. When TbAUK3 was RNAi depleted at 24 hrs, we detected a modest reduction in the percentage total of EdU positive cells (∼80 % to ∼ 70 %), while at 48 hrs post induction the number of EdU positive cells was dramatically reduced (only ∼14% of cells), indicating that loss of TbAUK3 results in reduced DNA synthesis. We found no difference in EdU incorporation in parental 2T1 control cells in the absence or presence of tet (**Fig. S2I**). When combined with our cell cycle analyses (**Fig. S2G**), we suggest that the modest depletion of EdU positive cells at 24 hrs suggests *T. brucei* re-enters new rounds of DNA replication despite incomplete nuclear segregation, further supporting the lack of a canonical SAC. At 48 hrs, the number of cells in S-phase reduces from ∼20 % to ∼ 14 %. This, combined with the accumulation of zoid cells that lack nuclei, may suggest a close association between impaired nuclear genome segregation and DNA replication.

Increasing data suggests aurora kinases directly regulate DNA repair proteins, acting as putative modulators of the DNA Damage Response (DDR) [32]. To ask if TbAUK3 depletion correlates with enhanced DNA damage, we performed immunoblotting to test for increased phosphorylation of the core histone H2A, generating γH2A, a well-defined marker of DNA damage in *T. brucei* [33]. We induced RNAi of TbAUK3, then prepared total protein extracts at 24 and 48 hrs post induction. Proteins were then separated by SDS-PAGE electrophoresis and transferred to a PVDF membrane **(Fig. 2A&B; Fig. S3A**). Membranes were probed with anti-γH2A antiserum and anti-EF1A (as a loading control). γH2A signal was then normalized to the loading control and a fold-change in signal was calculated relative to uninduced controls at each timepoint. At 24 hrs post induction, we detected no significant increase in yH2A signal (∼1-fold), but at 48 hrs, yH2A signal was ∼ 3-fold increased, suggesting prolonged depletion of TbAUK3 results in DNA damage. To further test if the increase in yH2A signal indicates increased DNA damage, we examined by IFA the presence of RAD51 foci using *T. brucei* specific anti-RAD51 antiserum. RAD51 is a key enzyme in homologous recombination (HR) that relocalises into nuclear foci that form at sites of damaged DNA in mammalian [34] and trypanosome cells[35]. In uninduced cells, we observed only ∼ 1% of cells in the population with nuclear RAD51 foci, but upon depletion of TbAUK3, the number of foci increased to ∼ 9 % (36 hrs) and ∼ 6 % (48 hrs) (**Fig. S3B&C**). Taken together with the cell cycle analysis, loss of TbAUK3 is associated with DNA damage and instability. Given a putative role for TbAUK3 in maintaining genome stability, we tested if TbAUK3 depletion enhanced sensitivity to damaging agents. We examined the parasite’s response to two inducers of DNA damage: Methyl Methane<u>s</u>ulfonate (MMS) and Ultra<u>v</u>iolet (UV) light. For MMS, we induced RNAi by adding tet then grew *T. brucei* in the absence and presence of 0.0003 % MMS. Every 24 hrs, cell density was assessed by counting. As expected, TbAUK3 depletion in the absence of damage reduced parasite proliferation. Exposure to MMS reduced proliferation in uninduced cells relative to untreated controls; however, upon TbAUK3 depletion, *T. brucei* became significantly sensitized to MMS (**Fig. 2C**). To test for sensitivity to UV, TbAUK3 RNAi was induced for 24 hrs by the addition of tet then at 24 hrs post induction, cells were exposed once to 1500 J/m^2^ UV. Cellular density was then assessed 24 hrs and 48 hrs post UV exposure (**Fig. 2D**). As expected, for unirradiated cells, parasite proliferation was reduced by RNAi relative to uninduced controls. After UV exposure, we found no significant differences in cell density or survival post exposure across all time points relative to uninduced (**Fig. 2D**). Taken together, TbAUK3 depletion enhances DNA instability and sensitivity to MMS but not UV radiation, indicating a specific and not generalised role in DNA damage repair.

**Fig. 2.**
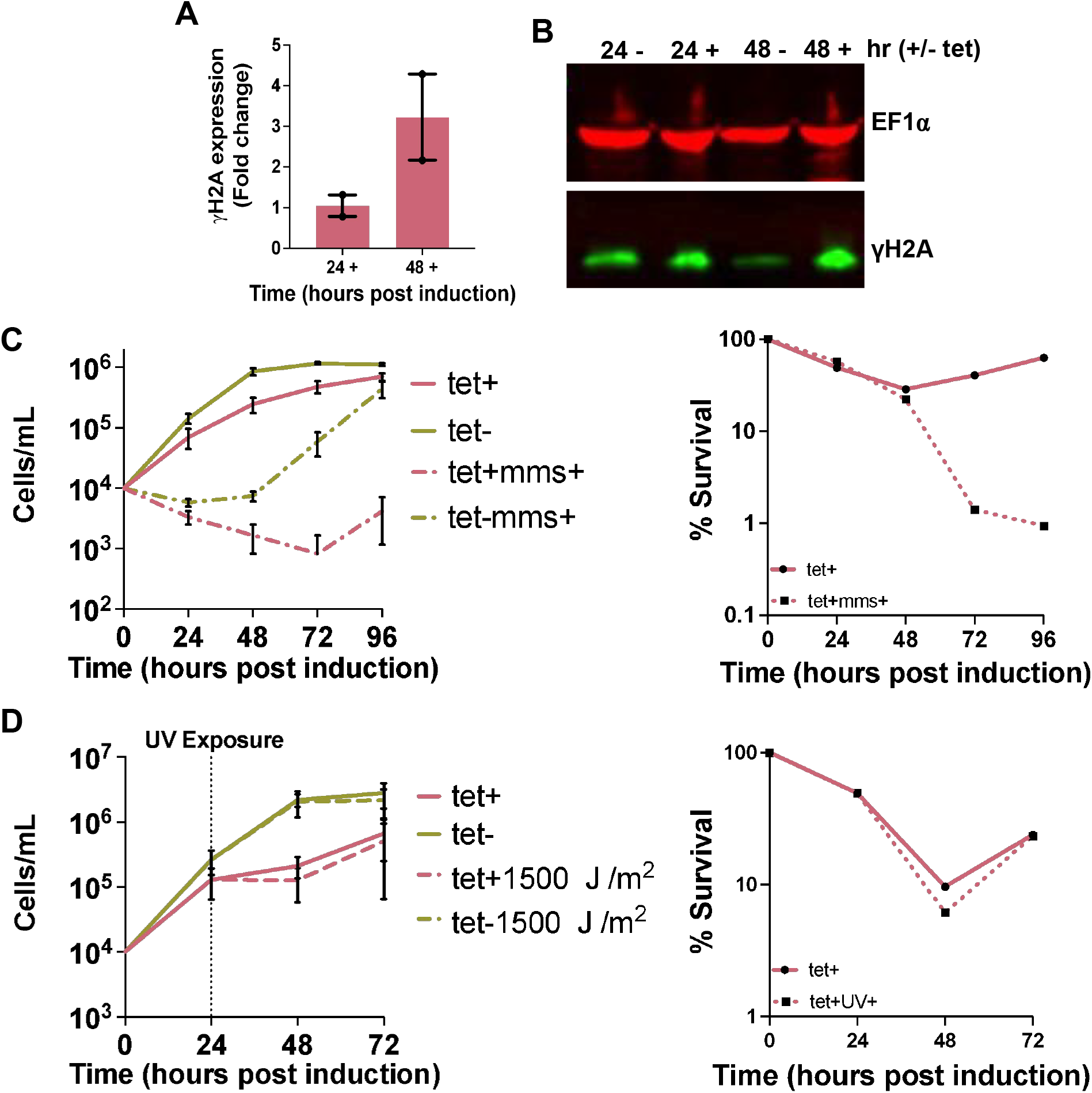
Depletion of TbAUK3 is associated with increased DNA damage and sensitivity to MMS but not UV exposure. (A) Immunoblotting analysis was performed to examine levels of γH2A in the absence and presence of tet induced TbAUK3 depletion. Signal for both proteins was quantified then γH2A signal normalized to EF1-alpha then a fold-change calculated relative to uninduced controls at 24 and 48 hrs respectively. Error bars = +/-SEM, n = 2 independent experiments. (B) Representative immunoblot image. *T. brucei* specific anti-γH2A antiserum was used at a concentration of 1:1000 and anti-EF1-alpha (Millipore) at a concentration of 1: 25,000 diluted in 5 % milk 1x PBS-Tween. Protein sizes were confirmed by loading Chameleon Duo Pre-Stained Protein Ladder (Li-Cor). Secondary antibodies used: Goat anti-rabbit (1:10,000) IRDye800 and goat anti-mouse (1:10,000) IRDye600. Images were captured on an Odyssey CLx Imager (LiCor). Proteins were separated by SDS-Page electrophoresis using 12 % Bis-Tris Novex PreCast^(R)^ cells run in a 1x MOPS buffer as per the manufacturer’s instructions. (C) To test sensitivity to MMS, cells were seeded at 1 ×10^4^ cells. mL^-1^ then RNAi induced with the addition of tet to the medium. A stock of 0.3 % MMS was prepared in HMI-11 and was then diluted to the appropriate concentration. Cell density was assessed by counting using a Neubauer chamber every 24 hrs for 96 hrs. Right graph shows the percentage survival calculated relative to the uninduced controls. Error bars = +/-SEM, n = 3 independent experiments. Survival curves show the average values of the three experiments. (D) To test sensitivity to UV, cells were seeded at 1 ×10^4^ cells. mL^-1^ then RNAi induced with the addition of tet to the medium. After 24 hrs induction, cells were then exposed once to UV using a Stratalinker 2400 (Stratagene) at the stated dose. Cell density was assessed by counting using a Neubauer chamber every 24 hrs for 48 hrs total post UV exposure. Right graph shows the percentage survival calculated relative to the uninduced controls. Error bars = +/-SEM, n = 3 independent experiments. Survival curves show the average values of the three experiments.

In summary, we have shown that depletion of TbAUK3 is associated with reduced parasite proliferation in *in vitro* cultures of BSF *T. brucei*. We detected a wide range of nuclear segregation defects, including multi-nucleated cells and cells devoid of any apparent nuclear DNA. Moreover, we have shown that TbAUK3 depletion induces nuclear genome instability and damage in *T. brucei*. Together with the prior report of TbAUK3 localisation at the BB in PCF cells, our results are consistent with roles for this kinase during chromosome segregation and the DDR in these divergent eukaryotes. Further study is now required to describe these roles in greater detail, including assessing spindle assembly formation directly in *T. brucei* BSF cells in the absence of TbAUK3.

## FIGURE LEGENDS - SUPPLEMENTARY

**Fig. S1.**
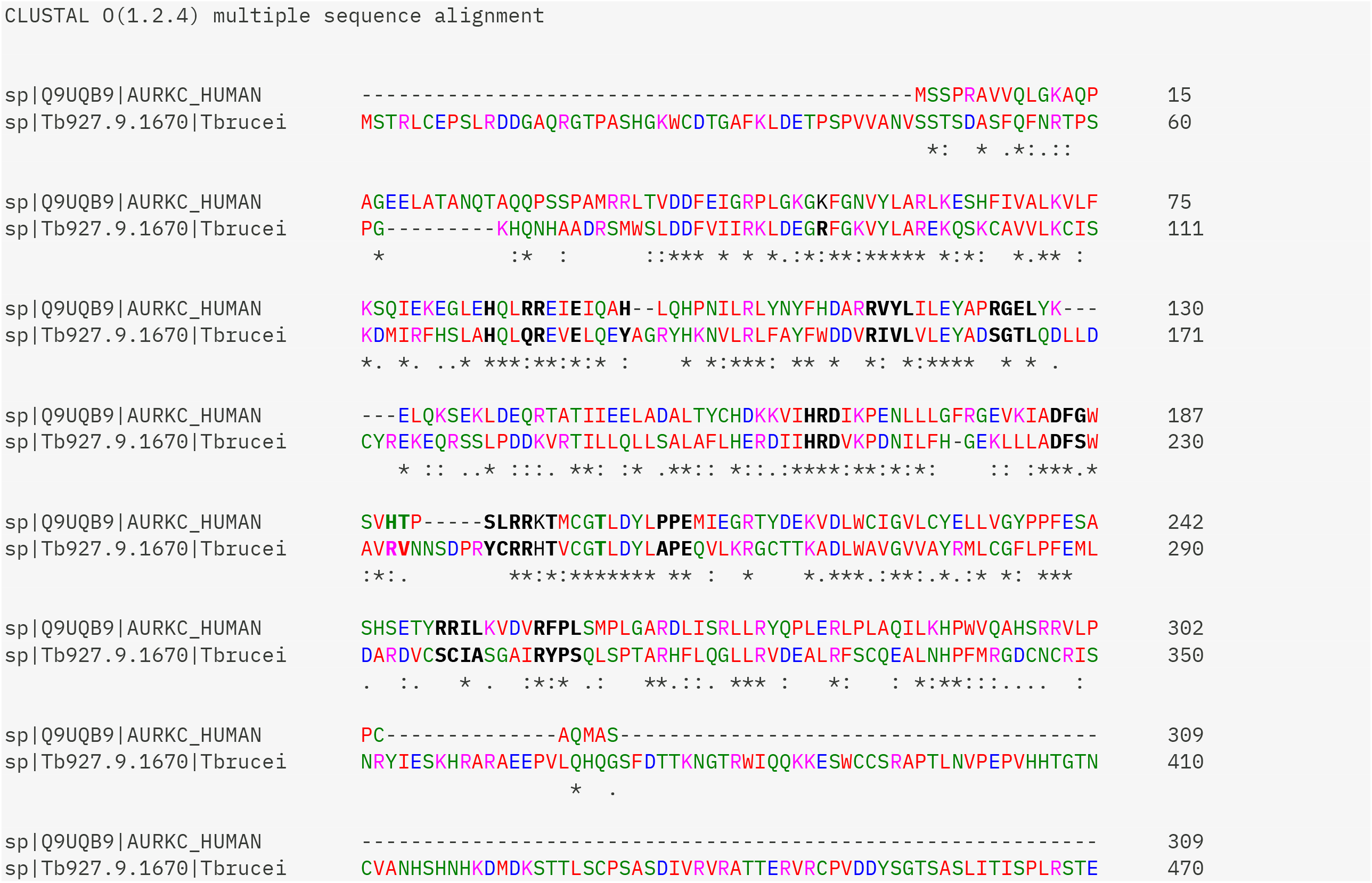
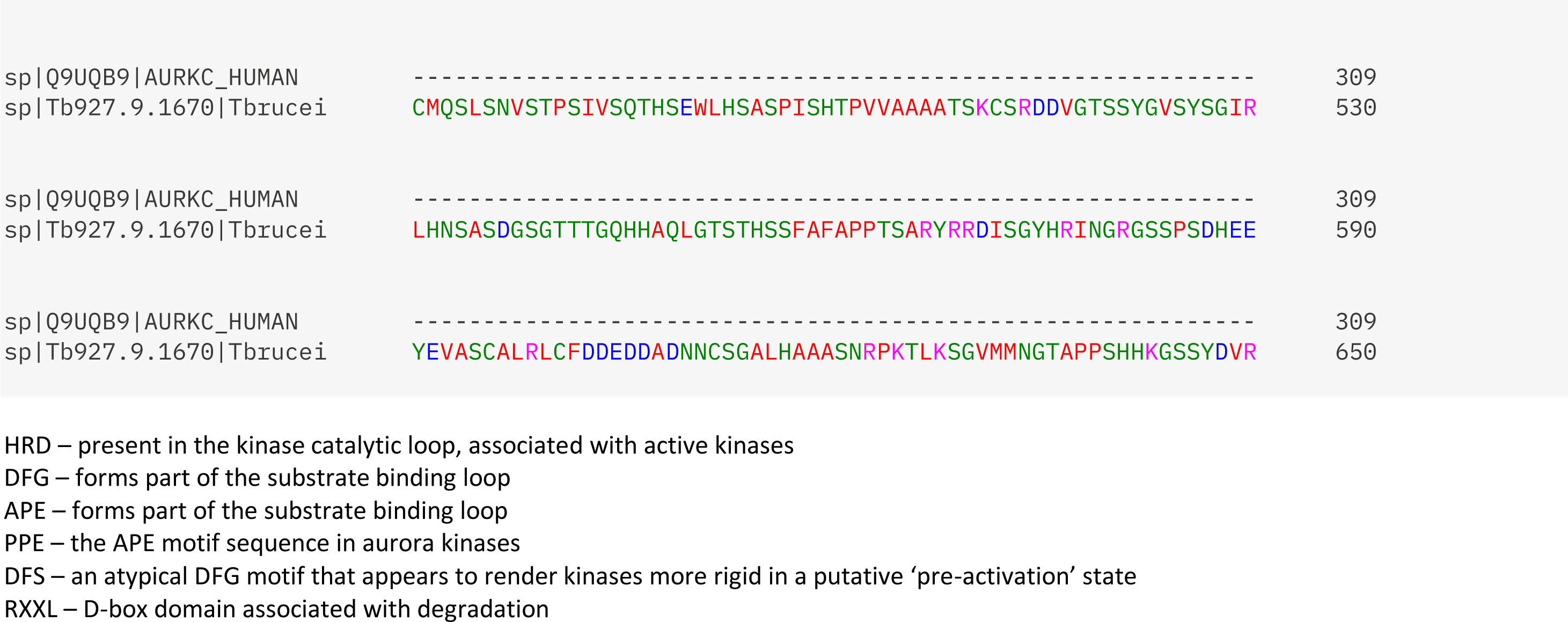
Amino acid sequence alignment of TbAUK3 to HsAURKC. The amino acid sequence of TbAUK3 (Tb927.9.1670) was aligned to the amino acid sequence of HsAURKC (QU9UQB9) using ClustalOmega (1.2.4). Important domains are highlighted in bold.

**Fig. S2.**
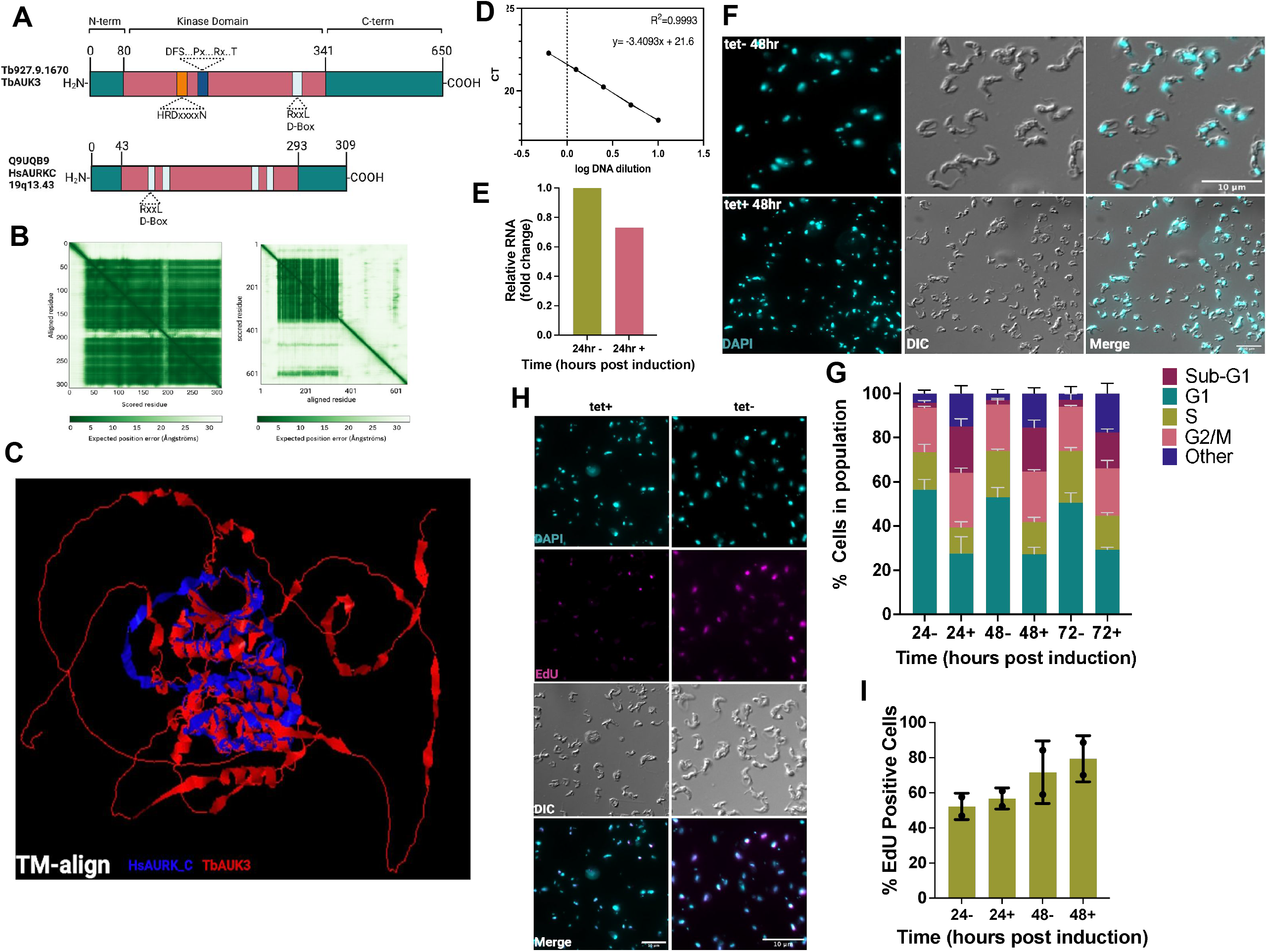
Depletion of TbAUK3 in BSF *T. brucei* coincides with reduced proliferation, nuclear segregation defects and reduced EdU incorporation. (A) Schematic diagram illustrating features of TbAUK3 compared with the human counterpart HsAURKC (B) Predicted Aligned Error (pAE) plot retrieved from the Alphafold 2 predictions of HsAURKC (right) and TbAUK3 (left) (C) Predicted overlap of TbAUK3 predicted protein structure with HsAURKC using TM align. Blue = HsAURKC, Red = TbAUK3 (D) Primer efficiency curve for primers used to test knockdown of TbAUK3. Genomic DNA was used to confirm efficiency (E) qRT-PCR test for knockdown of the mRNA of TbAUK3. RNA was extracted at 24 hrs in the presence and absence of tet induction using the Qiagen RNeasy kit. One microgram of RNA was converted to cDNA (random hexamers). Reverse transcriptase negative controls were run for each sample. Actin was used as an endogenous control gene for normalization. n = 1 experiment performed as technical triplicates. (F) Field of view images of cells in the absence (top panel) and presence (lower panel) tet induction at 48 hrs. DAPI (cyan). Scale bar = 10 µm. Images were captured on an Axioskop 2 63x lens. Brightness and contrast were altered in ImageJ to improve visualization. (G) Percentage of each cell cycle stage quantified by flow cytometry. Approximately 2.5 ×10^6^ cells were fixed in 1% formaldehyde for 10 mins at room temperature then washed, quenched in glycine (2.5 M), permeabilized in triton-X (0.01 % in PBS) then treated with 100 µg. mL^-1^ RNaseA to digest RNA (for 30 mins at 37 °C). DNA was stained using Propidium Iodide (PI; 10 µg. mL^-1^). Samples were filtered using a nylon membrane then events captured on an BD Calibur using the PE-CF594 filter for PI detection. Representative FAC plots and gating strategy are available in Supplementary Data 4. Error bars = +/-SD, n = 3 independent experiments. (H) Representative field of view images of EdU incorporation in the absence (left) and presence (right) of tet induction at 48 hrs. DAPI (cyan), EdU (magenta) Scale bar = 10 µm. Images were captured on an Axioskop 2 63x lens. Brightness and contrast were altered in ImageJ to improve visualization (I) The total number of cells in the population positive for nuclear EdU signal in 2T1 control cells, expressed as a percentage of the total number of cells counted. Error bars = +/-SD, n = 2 independent experiments, > 100 cells were counted/experiment.

**Fig. S3.**
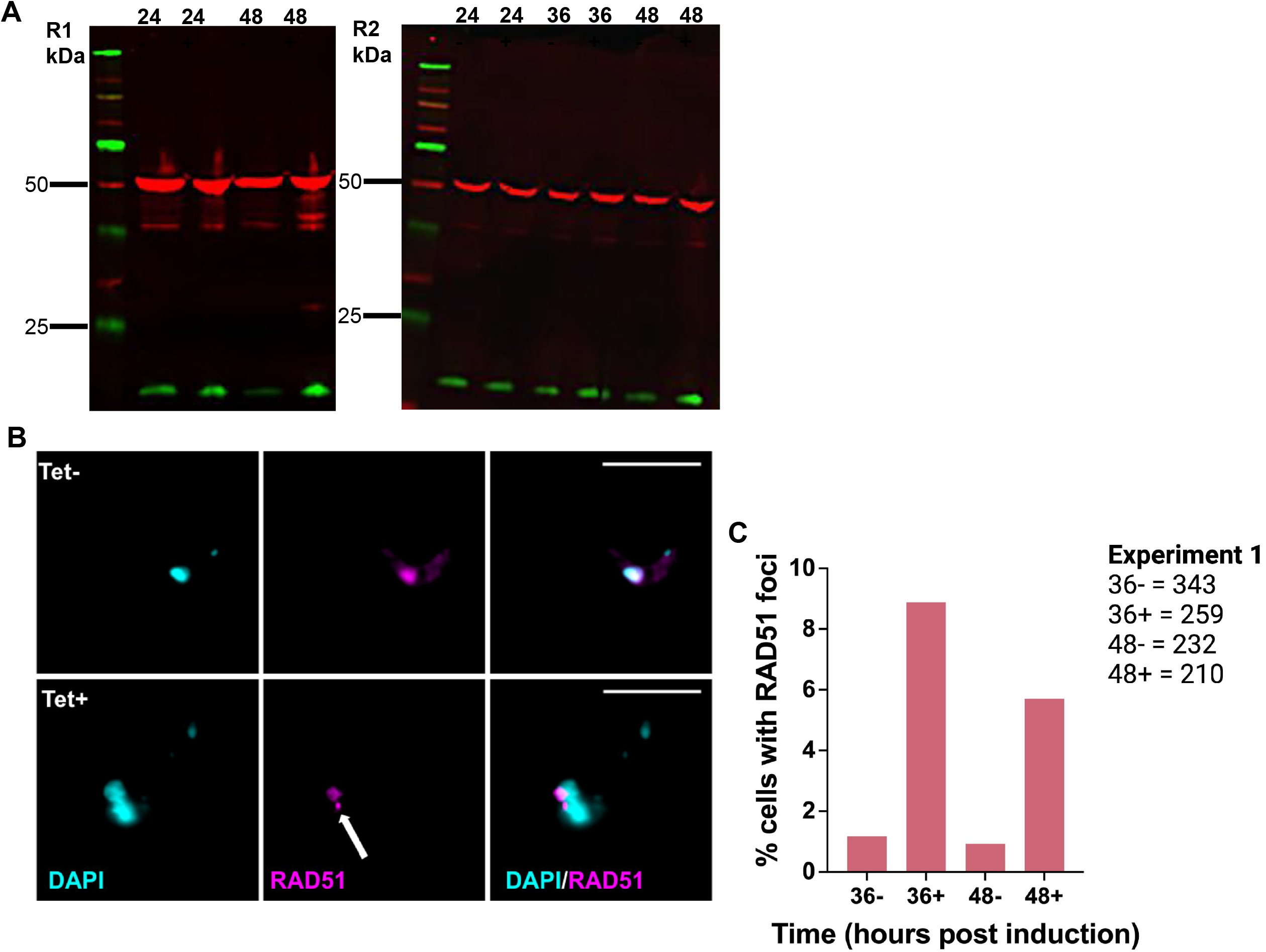
RAD51 foci accumulate in TbAUK3 depleted BSF cells. (A) Immunoblots relating to Figure 2A&B. (B) Representative images of RAD51 signal in the absence (top) and presence (lower) of tet induction at 48 hrs. DAPI (cyan), RAD51 (magenta). White arrow indicated a RAD51 foci. Scale bar = 10 µm. Images were captured on an Axioskop 2 63x lens. Brightness and contrast were altered in ImageJ to improve visualization. (C) Graph shows the number of RAD51 foci positive nuclei as a percentage of the total number of cells counted in the population. Number of cells counted is shown beside the graph. n = 1 experiment.

## AUTHOR CONTRIBUTIONS

JAB planned experiments, performed experiments, collected, and analyzed data, and wrote the manuscript. BP and AI performed experiments for Figure 1 and Figure 2 and collected data. BG wrote and critically reviewed the manuscript. LROT and RM critically revised the manuscript. RM additionally secured funding for the study. Figures were prepared using MS Powerpoint.

## ACKNOWLEDGEMENTS

This work was supported by the Wellcome Trust [224501/Z/21/Z], the BBSRC [BB/N016165/1, BB/R017166/1, BB/W001101/1], the MRC [MR/S019472/1], and FAPESP [2024/08412-0; JAB] & [23/03015-0; LROT]. The Wellcome Centre for Integrative Parasitology was supported by core funding from the Wellcome Trust [104111].

## APPENDIX A

## SUPPLEMENTARY DATA

The following data are supplementary to this article:

Supplementary_Figure 1

Supplementary_Figure 2

Supplementary_Figure 3

Excel sheet containing raw data underlying figures

